# Adenosine Kinase regulates Sleep Timing and the Homeostatic Sleep Response through Distinct Molecular Pathways

**DOI:** 10.1101/2023.12.05.570070

**Authors:** Zeinab Wakaf, Quang Dang, Yining Ru, Lewis Taylor, Sejal Kapoor, idhar Vasudevan, Robert Dallmann, Aarti Jagannath

**Author notes:** Lead contact AJ.

## Abstract

Sleep behaviour is broadly regulated by two drives, the circadian (Process C), which is orchestrated by the suprachiasmatic nuclei (SCN), and controls sleep timing, and the homeostatic (Process S), which controls sleep amount and the response to sleep deprivation (Borbély *et al*., 2016). However, the molecular pathways that mediate their independent effects, and their interactions remain unclear. Adenosine is an important integrator of both processes (Bjorness & Greene, 2009; Jagannath *et al*., 2021, 2022), such that adenosine levels track and modulate wakefulness, whilst adenosine signalling inhibits the circadian response to light. Therefore, we studied the sleep/circadian behaviour, and cortical and SCN transcriptomic profiles of a mouse model overexpressing Adenosine Kinase (Adk-Tg) (Fedele *et al*., 2005), (Palchykova *et al*., 2010). We found that overall, the Adk-Tg mouse slept less and showed lower amplitude circadian rhythms with an altered sleep/wake distribution across the 24h day, which correlated with changes in transcription of synaptic signalling genes that would shift the excitatory/inhibitory balance. In addition, the Adk-Tg mouse showed a reduced level of ERK phosphorylation, and attenuation of DNA repair related pathways. After sleep deprivation, however, the Adk-Tg mouse significantly increased relative to wildtype, immediate early gene expression levels including of *Arc*, but paradoxically reduced ERK phosphorylation. Thus, baseline sleep levels and timing are regulated by ERK signalling, whereas the response to sleep loss is mediated by the alteration of the transcriptomic landscape independently of ERK.

## Introduction

Sleep is a complex global state that involves changes in consciousness, brain activity and motor control and in rodents, broadly consists of two phases, non-rapid eye movement (NREM) sleep and rapid eye movement (REM) sleep, that alternate cyclically across sleep. NREM sleep, or slow wave sleep, is defined by low frequency high amplitude EEG waves and REM sleep, known as active sleep, is characterized by high-frequency low amplitude waves, rapid eye movement and muscle atonia. Although sleep is a biological state observed in almost all organisms, our understanding of the molecular mechanisms that regulate sleep and sleep need remain poor.

The sleep/wake transition is broadly controlled by two major drives; the homeostatic - driven by sleep pressure or sleep need, which correlates with prior wake, also known as Process S, and circadian Process C (Borbély *et al*., 2016). The latter is driven by a well characterised cell autonomous molecular clock aligned to the environmental light dark cycle by a master clock in the suprachiasmatic nuclei (SCN) (Hughes *et al*., 2015). Process S has two distinct but related components – baseline sleep and homeostatic responses. The latter is defined by the amount and depth (the quantification of delta power or slow-wave activity – SWA) of recovery sleep following sleep deprivation. These drives were first described over 4 decades ago, yet how they interact to shape sleep/wake remains unclear. This is primarily as the molecular signals used by the homeostatic drive remain to be definitively identified. Moreover, if and where these molecular signals converge with the molecular circadian clock to ultimately influence sleep/wake behaviour are poorly understood.

Genetic studies have identified a role for kinases, including ERK1/2, SIK3 and CaMKIIα/β in regulating some molecular correlates of Process S. For instance, loss of *Erk1* and *Erk2* in cortical neurones increases the duration of wakefulness in mice, while their pharmacological inhibition reduces expression of *Arc* and *Homer1a* in neuronal cultures, two typical genes associated with extended wakefulness (Cirelli *et al*., 2004a; Maret *et al*., 2007; Mikhail *et al*., 2017). Similarly, gain-of-function allele of *Sik3* in excitatory neurones, mainly glutamatergic, increases NREM delta power and NREM amount, that correlate with sleep need and sleep depth, respectively (Funato *et al*., 2016; Kim *et al*., 2022). Additionally, embryonic knockout of *Camk2α* or *Camk2*β and post-natal inhibition of CAMKII both result in significant decrease in sleep duration (Tatsuki *et al*., 2016; Tone *et al*., 2022). In addition, studies have profiled the transcriptomic changes associated with homeostatic response to sleep deprivation. The genes consistently reported as being upregulated upon sleep deprivation include *Homer1*, *Activity regulated cytoskeleton-associated protein* (*Arc),* immediate early genes (IEGs) and *Brain-derived neurotrophic factor* (BDNF). The circadian clock genes are also regulated by sleep deprivation, such that *Per1*/2 are upregulated, whilst *Dbp* is downregulated (Cirelli *et al*., 2004b; Terao *et al*., 2006; Mackiewicz *et al*., 2007; Wisor *et al*., 2008; Thompson *et al*., 2010a; Hinard *et al*., 2012; Gerstner *et al*., 2016). Similar transcriptomic changes were also evident in *in vitro* cortical cultures (Hinard *et al*., 2012). Nevertheless, upstream regulation of these molecular pathways, and their precise roles in the different aspects of sleep regulation, namely sleep timing, the distribution of sleep/wake episodes across the 24h day, and the response to sleep loss are poorly understood.

Since adenosine is an important signal to both Process S and Process C (Bjorness & Greene, 2009; Jagannath *et al*., 2021, 2022), the adenosine signalling pathway represents a target by which to disrupt the relationship between different aspects of sleep regulation and probe the underlying molecular changes. Extracellular adenosine levels, as a by-product of ATP and cAMP, reflect neuronal activity, regulate immediate early gene transcription and signal Process S through adenosine A1 and A2A receptors (Basheer *et al*., 1999; Elmenhorst *et al*., 2007; Brown *et al*., 2012; Lazarus *et al*., 2019). Adenosine agonists have been shown to enhance SWA, while adenosine receptor antagonists like caffeine promote wakefulness (Bjorness & Greene, 2009). Recent studies show that adenosine signalling is one of the molecular pathways by which Process S and Process C interact, allowing sleep/wake history to modulate circadian entrainment (Jagannath *et al*., 2021, 2022). Caffeine, whose effects on wakefulness are mediated by adenosine receptors (Lazarus *et al*., 2011) is known to increase photic phase-shifts of the circadian system in both humans and mice and lengthen circadian period, whilst sleep deprivation reduces the effects of light on the clock (Challet *et al*., 2001; Oike *et al*., 2011; van Diepen *et al*., 2014; Burke *et al*., 2015; Ruby *et al*., 2018). Furthermore, adenosine receptor antagonists can both phase-shift the clock and mediate entrainment (Antle *et al*., 2001). *In vivo* manipulations of adenosine kinase (ADK), the primary metabolising enzyme regulating extracellular adenosine levels (Boison, 2006), have further shed light on the role of adenosine in both Process S and C. Mice overexpressing ADK (Adk-Tg) exhibit reduced adenosinergic tone in the brain (Fedele *et al*., 2005). Interestingly, Adk-Tg mice exhibit less SWA build up, but paradoxically following sleep deprivation, they have higher rebound NREM sleep than WT (Palchykova *et al*., 2010). This suggests that the baseline sleep amount and timing could be regulated separately from the homeostatic sleep response, in a manner linked to divergent pathways downstream of adenosine. Furthermore, Adk-Tg mice show enhanced circadian phase-shifting response to light and the molecular pathway by which adenosine signalling regulates the clock involves the sleep-regulating kinase ERK1/2 (Jagannath *et al*., 2021). Hence, this animal provides an excellent model in which to investigate the interaction between Process C and S, and the kinase/transcriptomic molecular correlates of the different aspects of sleep regulation.

In this paper, we examine the sleep/circadian behavioural phenotype of the Adk-Tg mouse and describe the downstream transcriptomic response of the SCN and the cortex under basal conditions and following sleep deprivation. We show that in addition to sleeping less overall, the Adk-Tg mouse displays an altered distribution of sleep across the 24h light-dark cycle. At the transcriptomic level, we observe altered expression of GABAergic and glutamatergic signalling components within both the SCN and cortex, in addition to changes in oxidative phosphorylation, DNA repair and purine metabolic pathways. Immediate early gene expression in Adk-Tg mice is lower in the cortex which could be explained by lower levels of ERK1/2 and ERK1/2 phosphorylation (pERK1/2). We show that ERK phosphorylation status correlates with the amount of sleep under baseline conditions such that the Adk-Tg mouse, which shows reduced sleep also shows lower levels of pERK1/2. However, pERK1/2 levels are strikingly reduced following sleep deprivation in both WT and Adk-Tg mice, which is unexpected given pERK1/2 levels typically increase with wakefulness, and therefore ERK signalling cannot underlie rebound sleep. In contrast, we found that *Arc* and IEG expression are increased relative to WT in the Adk-Tg after sleep deprivation, which could explain the increase in rebound NREM sleep despite lower pERK1/2 following sleep deprivation in the Adk-Tg.

## Results

### Adk-Tg mice display altered distribution of sleep across the 24h light-dark cycle

In order to investigate the effects of extracellular adenosine reduction on sleep and circadian behaviour, we recorded activity using passive infrared receivers (PIRs), from which inactivity levels have been well validated as a non-invasive measure of sleep (Fisher *et al*., 2012). Under normal environmental conditions or 12-hour light-dark (LD) cycles, Adk-Tg mice display more light-phase activity (Figure 1A-C) relative to WT, with significantly higher interdaily variability (Figure 1D, p-value 0.019) and lower intradaily stability (Figure 1E, p-value 0.0006). Moreover, Adk-Tg mice demonstrate a lower amplitude of activity (Figure 1F, p-value 0.04). Interestingly, the increased activity observed during the day is paralleled by a significant increase in immobility-defined sleep in the early hours of dark, and a significant decrease later during the night (Figure 1G), suggesting an altered distribution of sleep/wake patterns across the 24h day. Cumulatively however, Adk-Tg display more wakefulness than the WT throughout the light-period, as reported previously using EEG (Palchykova *et al*., 2010), while we found no overall difference during their active period or dark phase (Figure 1H).

**Figure 1:**
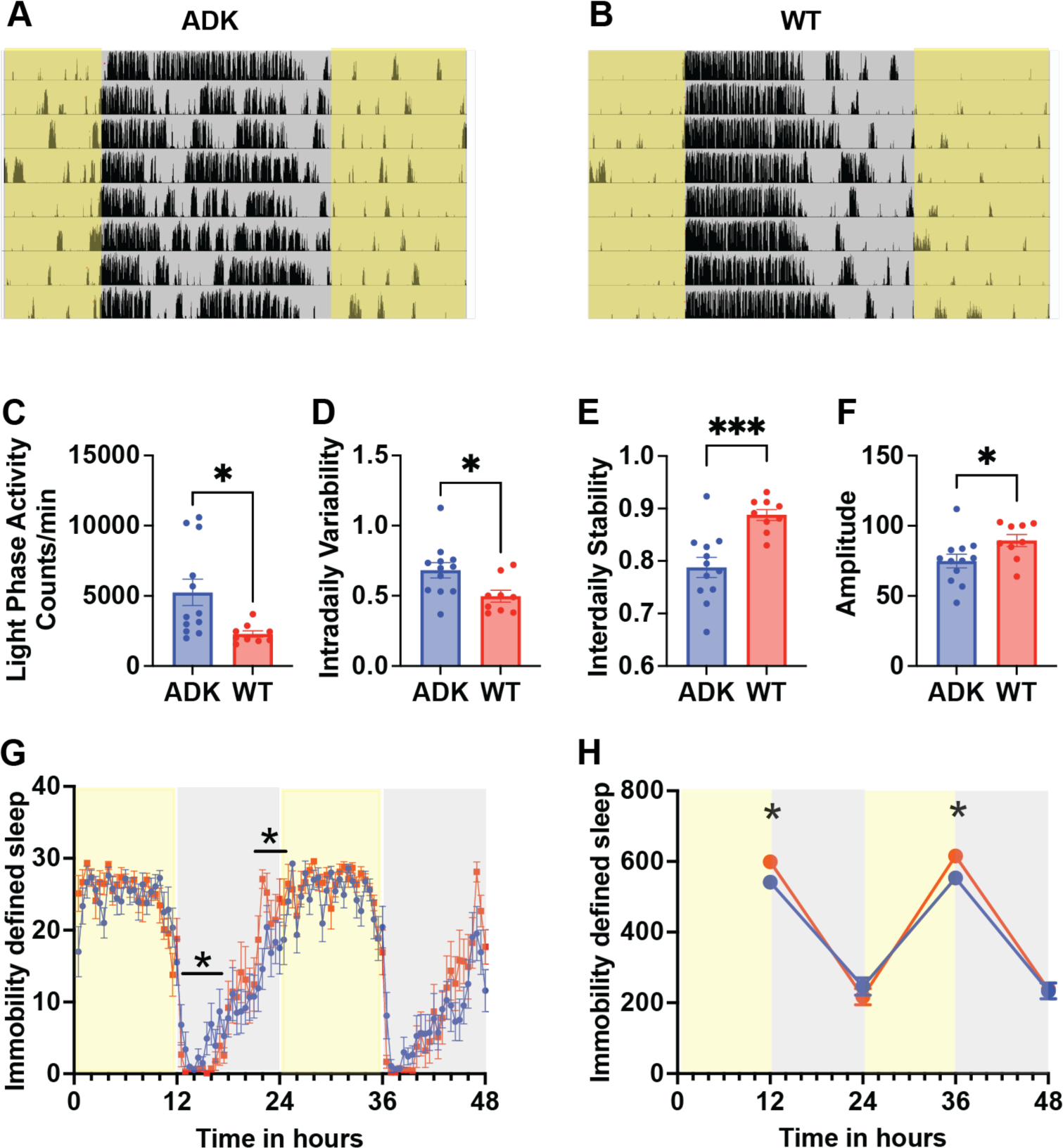
Adk-Tg mice show altered distribution of sleep and lower amplitude circadian rhythms under 12/12 LD cycles. Representative actograms from (A) Adk-Tg (ADK) and (B) wildtype (WT) mice under a 12/12 L/D cycle, yellow indicates lights on, grey indicates lights off and black bars are PIR indicated activity. Quantification of activity patterns showed significant differences between the genotypes in (C) activity during the light phase (D) intradaily variability (E) interdaily stability and (F) amplitude. Genotype differences in immobility defined sleep (in minutes) (G) at the start and end of the dark (active phase) and (H) in total sleep levels during the light (inactive phase) were detected. n=10 per genotype, * = p<0.05, *** = p<0.001 with a student’s t-test for C-F. * = p<0.05 with paired t-tests for G and * = p<0.05 with Sidak’s multiple test correction after two-way ANOVA analysis for H (F(3, 57) = 5.380 for time x genotype).

### Adk-Tg mice display altered distribution of sleep and altered circadian rhythmicity in constant dark

In constant dark (DD) Adk-Tg mice have a longer period (Figure 2A-C, p-value 0.0001), lower amplitude (Figure 2D, p-value 0.0004) and higher intradaily variability (Figure 2E, p-value <0.0001) relative to WT. The increase in sleep fragmentation in Adk-Tg mice becomes more apparent under DD, with Adk-Tg mice displaying significantly lower counts of activity but more bouts of activity throughout the day (Figure 2F, counts/bout p-value 0.0006, bouts/day p-value 0.0006) compared to WT. Moreover, like what is seen in LD, Adk-Tg mice sleep significantly more in the first hours of activity (Figure 2G, CT12-18, p-value <0.001) but sleep less in the second half of their activity period (Figure 2G, CT18-24, p-value <0.001). This difference seems to be more pronounced in DD than in LD. However, cumulatively across the day, Adk- Tg mice show no difference in immobility-define sleep, and in the absence of light there is no difference in sleep during their subjective inactive phase. Together these findings suggest circadian regulation of the sleep-mediating effects of pathways downstream of adenosine signalling during the active period, and an important role for the light-adenosine signalling axis in remodelling sleep timing.

**Figure 2:**
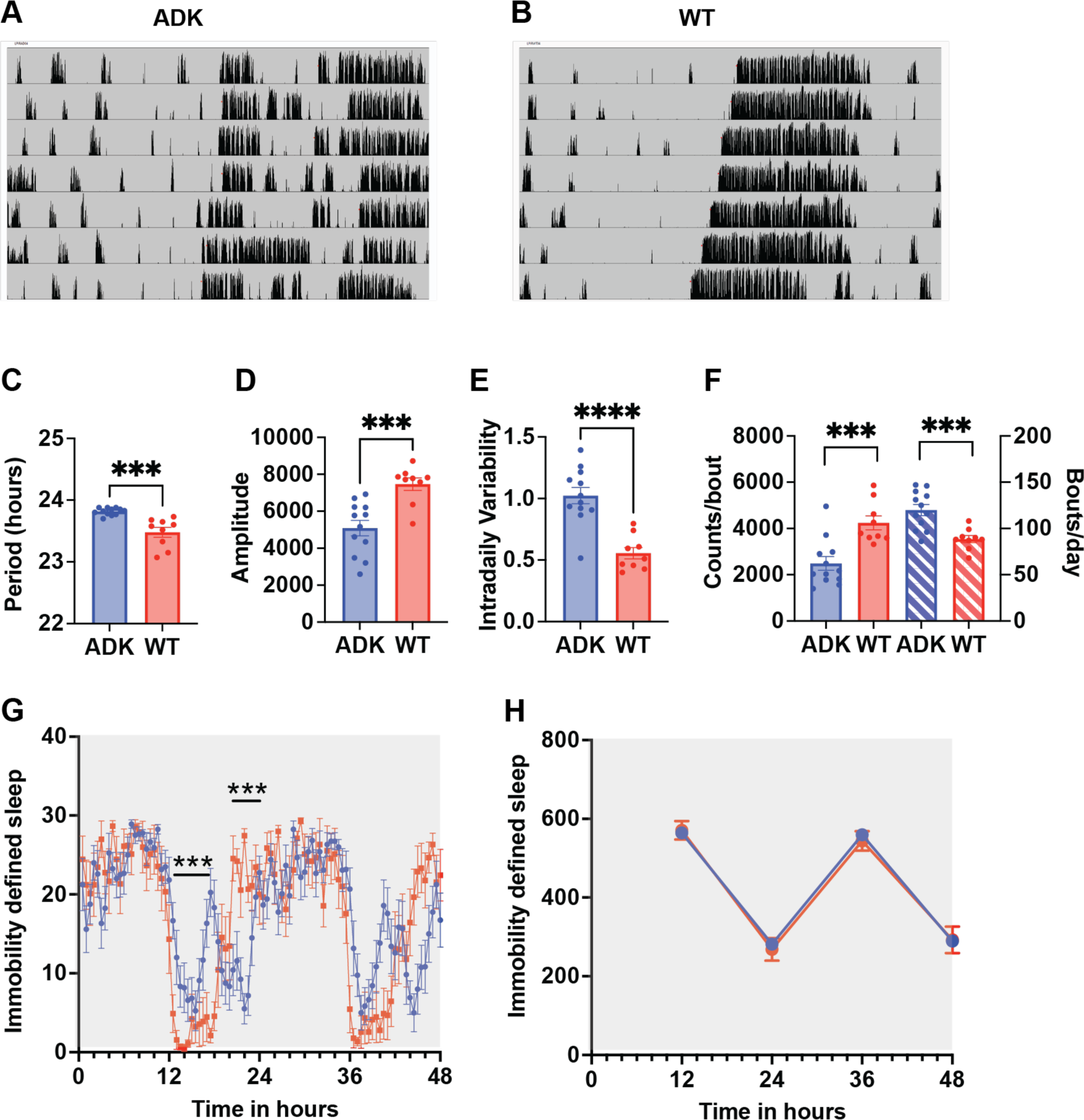
Adk-Tg mice show altered distribution of sleep and lower amplitude circadian rhythms under constant dark. Representative actograms from (A) Adk-Tg (ADK) and (B) wildtype (WT) mice under constant dark, grey indicates lights off and black bars are passive infrared indicated activity. Periodogram analysis showed significant differences between the genotypes in (C) circadian period (D) amplitude (E) intradaily variability and (F) counts/bout. Differences in immobility defined sleep (in minutes) (G) at the start and end of the dark (active phase) as plotted across 2 days in DD, but no overall differences (H) in total sleep levels were seen. n=10 per genotype, *** = p<0.001, **** = p<0.0001 with a student’s t-test for C-F. *** = p<0.001 with paired t-tests for G, Two-way ANOVA analysis for H showed no significant differences.

Similar to DD, in constant light (LL) Adk-Tg mice also exhibited increased fragmentation of sleep, with significantly higher number of sleep bouts across the day and a lower sleep count per bout of sleep (Supplementary Figure 1C and 1D). However, although Adk-Tg and WT mice both displayed similar distribution of immobility-defined sleep across the day under LL, Adk-Tg mice overall slept less than their WT controls (Supplementary Figure 1E and 1F), emphasizing the role of light in promoting the sleep phenotype in Adk-Tg. Furthermore, unlike in DD, there is no significant different in circadian period length (data not shown), again suggesting a difference in clock function where light input is modulated by adenosine levels.

### Adk-Tg mouse shows altered SCN and cortical transcriptional profiles

To assess the molecular pathways downstream of adenosine signalling mediating the sleep phenotypes, we performed transcriptomic analysis of the SCN and cortex for both genotypes under baseline 12:12 LD conditions (Figure 3). As expected, Adk-Tg mice expressed higher levels of ADK in both SCN and cortex (log2 fold change - L2FC 1.77 adj. p-value 6.9E-126 Figure 3A, and L2FC 1.89 adj. p-value 4.54E-218 3B, respectively). Similarly, gene ontology analysis of the transcriptome of both brain regions displayed an enrichment for ADK activity, mitochondria and cellular metabolism (Figure 3C, 3D), all of which are associated with adenosine regulation and ATP synthesis and degradation. This demonstrates the role of adenosine loss in the observed behavioural phenotypes and validates the mouse model used in this study.

**Figure 3:**
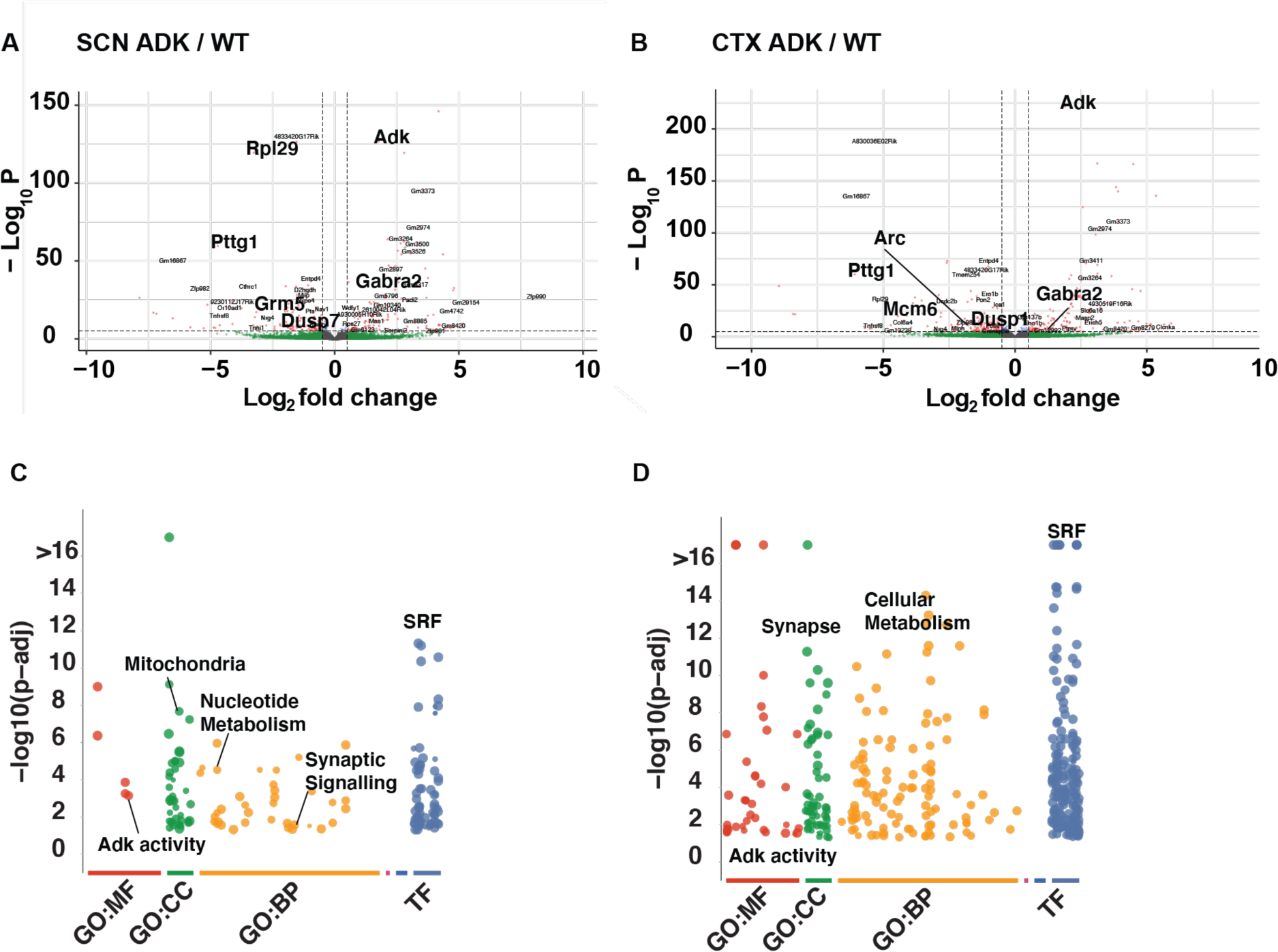
Adk-Tg mice show altered regulation of immediate early genes and synaptic signalling genes in SCN and Cortex. Volcano plots of RNA-Seq data from (A) SCN and (B) Cortex (CTX) of Adk-Tg (ADK) versus wildtype (WT) mice at CT6. Dotted line horizontal indicates adjusted -Log_10_ P = 5, dotted vertical lines indicate Log_2_ fold change of ±0.5. Green = transcripts with Log_2_ fold change > ±0.5, red = transcripts with Log_2_ fold change > +_ 0.5 and adjusted p < 0.0001. GO analysis for the significantly differential transcripts in SCN (C) and Cortex (D), indicating increased Adk activity as expected, but also altered nucleotide metabolism and synaptic function. (GO = Gene Ontology, MF = molecular function CC = cellular compartment, BP = biological process and TF = transcription factor prediction from Transfac.

At baseline conditions (sleep at CT6), the SCN and cortex behaved similarly to overexpression of ADK. Both brain regions displayed an increase in GABAergic signalling and downregulated glutamatergic signalling (Figure 3A, B). In addition, the SCN showed decreased expression of potassium inwardly rectifying channels including *Kcnj4* and *Kcnj13*, which together suggest an altered synaptic function. Furthermore, in line with increased ADK expression (Figure 3 A, B) the Adk-Tg mice showed increased nucleotide metabolism and ADK activity (Figure 3 C, D). Serum response factor (SRF) was identified as the upstream transcription factor mediating the differences between ADK and WT (Figure 3 C, D). Moreover, the cortex of Adk-Tg mice showed a significant reduction in *Arc* (Figure 3C), an immediate early gene involved in long-term synaptic plasticity, memory formation and synaptic downscaling (Korb & Finkbeiner, 2011).

In the cortex under baseline conditions, genes involved in DNA damage repair like *Pttg1* (L2FC -6.8, adj. p value 1.2E-57), *Pnkp* (L2FC -0.347, adj. p value 0.004), *Mcm6* (L2FC -1.97, adj. p value 1.8E-31), *Mcm8* (L2FC -0.512, adj. p value 0.008), *Polb* (L2FC -0.203, adj. p value 0.004), and *Dtx3l* (L2FC -0.512, adj. p value 0.01) were all downregulated in Adk-Tg mice relative to WT, suggesting an overall lower ability to respond to DNA damage (see Supplementary Table 1). Similar to cortex, *Pttg1* (L2FC -4.7 adj. p value 5.8E-57) and *Mcm6* (L2FC -1.97, adj. p value 1.8E-31) were also downregulated in the SCN, while *Rif1* gene that is involved in repair of double strand breaks following DNA damage was upregulated (L2FC 0.284, adj. p value 0.003, Supplementary Table 1), proposing possible SCN DNA damage in Adk-Tg mice relative to their WT littermates.

### Arc and other IEGs are significantly upregulated in Adk-Tg following Sleep Deprivation

Palchykova et al. showed distinct responses of Adk-Tg mice to sleep deprivation (SD). Following 6 hours of SD, both Adk-Tg and WT mice showed an increase in NREM sleep and reduction in waking time (as expected to recover after SD) and baseline differences in REM. However, Adk-Tg mice had a significantly higher NREM rebound-sleep after SD relative to the WT controls (Palchykova *et al*., 2010). Aiming to understand the molecular pathways underpinning this increase in NREM rebound sleep, we performed transcriptomic analysis on SCN and cortex for Adk-Tg and WT mice following 6 hours of sleep deprivation (Figure 4).

**Figure 4:**
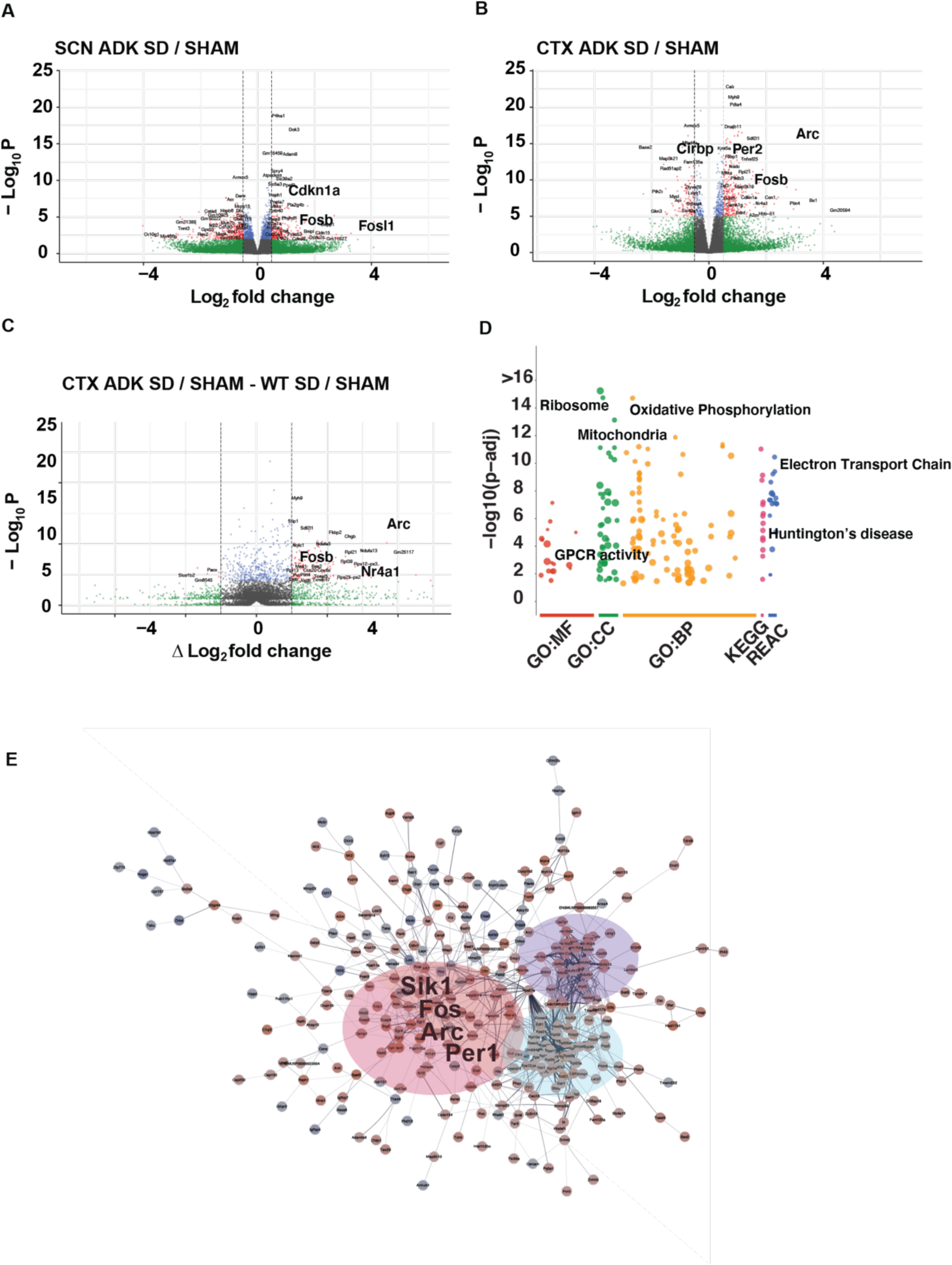
Adk-Tg mice show increased immediate early gene induction following sleep deprivation. Volcano plots of RNA-Seq data from (A) SCN and (B) Cortex (CTX) of Adk-Tg (ADK) sleep deprived (SD) for 6h (CT0-6) versus sham deprived (SHAM) mice. Relative difference in fold change between the ADK and WT responses to sleep deprivation in the Cortex (C). Dotted vertical lines indicate Log_2_ fold change of ±0.5. Green = transcripts with Log_2_ fold change > ±0.5, red = transcripts with Log_2_ fold change > +_ 0.5 and adjusted p < 0.01 (A), p<0.0001 (B) and p<0.01 (C). (D) GO analysis for the significantly differential transcripts in (C) indicating increased mitochondrial activity and translation in the ADK. (E) STRING analysis for the significantly differential transcripts in (C) indicating increased immediate gene induction (red bubble), oxidative phosphorylation / respiration (purple bubble) and ribosomal / translational machinery (blue bubble).

Following 6 hours of SD (Figure 4), as predicted, transcriptomes of SCN and cortex displayed upregulation of early response genes *like Fosb, Arc* and *Egr2* (Fig 4A, B, Supplementary Figure 2), and downregulation of *Dbp* and *Cirbp* in both genotypes (Figure 4). Furthermore, SD triggered an increase in DNA damage in both genotypes, as exemplified by an upregulation of *Cdkn1a* in SCN (Figure 4A, 4B, Supplementary Figure 2). Interestingly however, when comparing the relative fold-change differences between Adk-Tg and WT in response to SD, the upregulation of immediate early genes (IEGs) like *Arc, Fosb* and *Nr4a1* were significantly higher in the cortex of Adk-Tg mice (Figure 4C). This observation was further validated using qPCR, where relative mRNA expression of *Arc* and *Nr4a1* also showed a significant increase in both genotypes following SD (Supplementary figure 3), with a smaller baseline value for *Arc* in Adk-Tg mice and an enhanced upregulation following SD relative to WT. STRING analysis of the differentially upregulated transcripts also highlighted the prominent increase in immediate gene induction in Adk-Tg mice (Figure 4E). GO and STRING analyses also demonstrated significant increase in oxidative phosphorylation and electron transport chain processes in Adk-Tg mice, as well as an enhancement in ribosomal and mitochondrial functions (Figure 4D, 4E), which correspond to an increase in ATP-synthesis pathways likely compensating for the reduction in ATP and extracellular adenosine as a result of ADK overexpression (Kornberg & Pricer, 1951). The increase in oxidative phosphorylation in turn could be linked to a rise in ROS and compounding of DNA damage. Indeed, DNA damage/repair – related genes in response to SD were overexpressed in Adk-Tg mice after sleep deprivation. *Myc* (difference in log2 fold change - diffFC 0.518, adj. p-value 0.16) and the Growth arrest and DNA-damage inducible protein GADD45 gamma *Gadd45g* (diffFC 0.516, adj. p-value 0.1) that are involved in activating and responding to DNA-damage, were both upregulated (Supplementary table 1), and the DNA-damage sensor *Parp1* was also upregulated (diffFC 0.228, adj. p-value 0.07, Supplementary table 1). Moreover, the differentially upregulated transcripts in Adk-Tg mice following SD correlated with Huntington’s disease (Figure 4D), for which DNA repair mechanisms have been identified as key disease modifying pathways (Consortium, 2015). Together, these findings demonstrate that Adk-Tg display enhanced IEG induction following SD and would also exhibit attenuated DNA-repair abilities compared to WT, which could exacerbate SD-mediated DNA damage.

### ERK phosphorylation marks increased wakefulness in Adk-Tg, but does not correlate with “sleep need” following sleep deprivation in Adk-Tg or WT mice

The extracellular signal-regulated kinase 1/2 (ERK) is a member of the mitogen-activated kinase (MAPK) family and has been previously reported as a sleep-inducing kinase (Mikhail *et al*., 2017). Similar to Adk-Tg mice that display reduced sleep (Figure 1, 2), loss of *Erk1* and *Erk2* in cortical neurones has been shown to increase the duration of wakefulness, as does the inhibition of ERK phosphorylation. As an important factor in adenosine signalling (Shen *et al*., 2005), we set out to investigate ERK expression and phosphorylation in the Adk-Tg sleep phenotypes. Under sham conditions (sleep at CT6), there were ERK1 (*Mapk3*) or ERK2 (*Mapk1*) were not differentially expressed (Supplementary table 1). However, phosphorylation of ERK1/2 was significantly lower in Adk-Tg mice compared to their littermate controls (Figure 5). The fact that Adk-Tg mice experience increased wakefulness (Figure 1, 2) as well as a reduction in pERK1/2 (Figure 5) supports the role of pERK1/2 as a sleep-promoting kinase under baseline conditions (Mikhail *et al*., 2017), where phosphorylated ERK has been reported to increase in cortex throughout wakefulness and its accumulation is needed to promote sleep.

**Figure 5:**
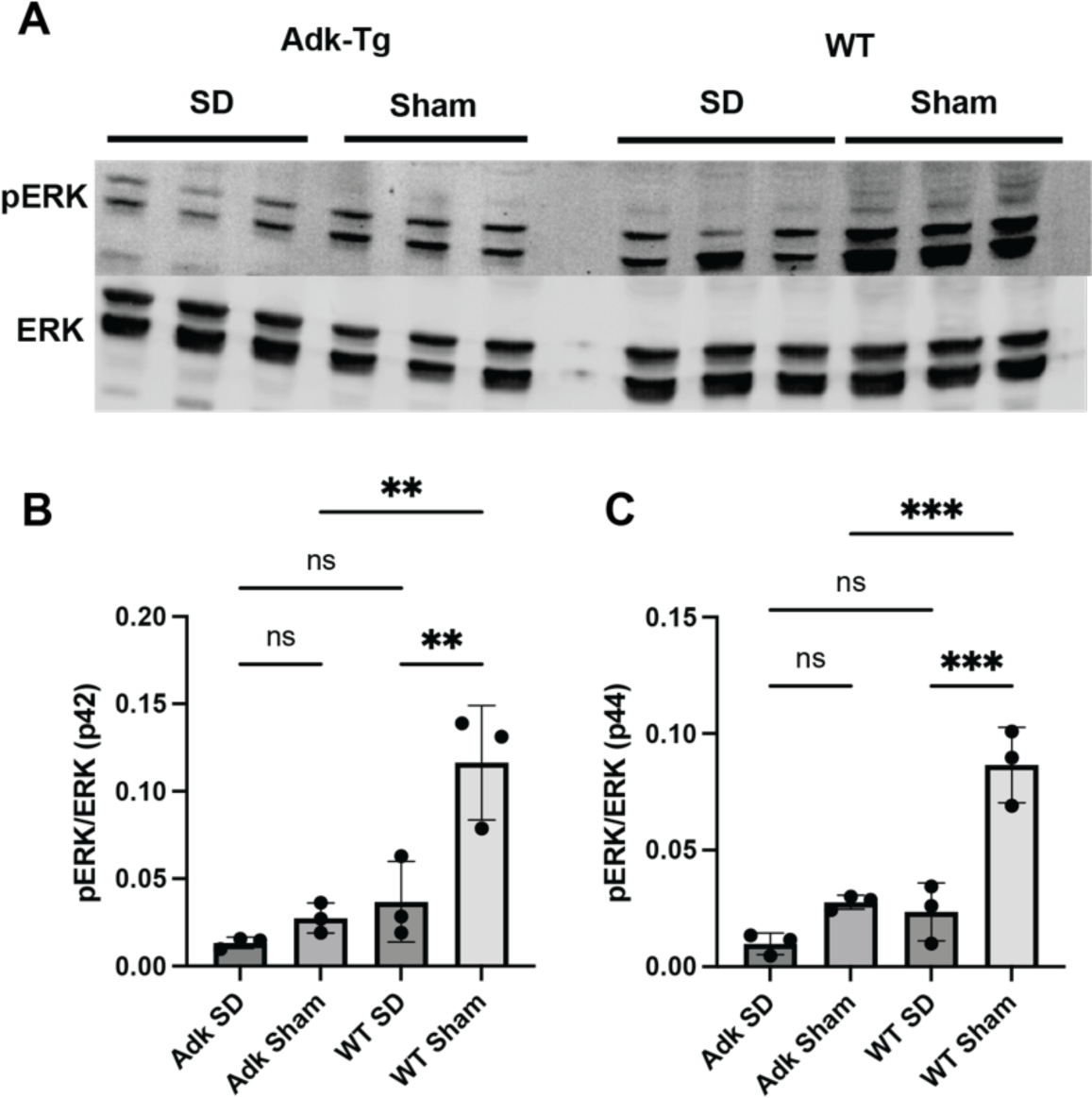
Adk-Tg mice show reduced ERK phosphorylation under baseline and sleep deprived conditions. (A) Western blots for ERK1/2 and phosphorylated ERK1/2 (pERK) in the cortex of Adk-Tg and wildtype (WT) mice sleep deprived (SD) for 6h (CT0- 6) versus sham deprived (Sham) mice. Bar charts (B) and (C) show quantification of pERK (p42 – ERK2 – B, and p44 – ERK1 – C) relative to ERK. n = 3, ** = p<0.01, *** = p<0.001 from Tukey’s multiple comparisons tests from one-way ANOVA analysis for B and C.

Following 6 hours of sleep deprivation, both genotypes exhibited reduction in pERK1/2 levels (Figure 5). Interestingly, SD has been shown to result in a significant increase in NREM rebound sleep in Adk-Tg mice relative to WT (Palchykova *et al*., 2010). Since pharmacological Inhibition of ERK-phosphorylation significantly reduces NREM rebound sleep following SD (Mikhail *et al*., 2017), our observations suggest ERK phosphorylation levels do not correlate with the increased rebound NREM phenotype of the Adk-Tg mouse under conditions of forced wakefulness, implying another regulatory pathway.

On the other hand, the upregulation of IEGs, particularly *Arc*, could be the key driver for rebound NREM sleep following SD in Adk-Tg mice. In *Arc* KO mice, the loss of *Arc* completely abolishes rebound upregulation of NREM and REM following SD (Suzuki *et al*., 2020), whilst the Adk-Tg mice show both increased *Arc* expression (Figure 4, Supplementary Figure 3) and increased rebound NREM sleep following SD (Palchykova *et al*., 2010). Hence, upregulation of *Arc* in ADK mice could be driving the rebound sleep. Indeed, immediate early genes known to be under the direct effect of *Arc* (such as *Fos*, *Egr1*, and *Nr4a1*) are upregulated in ADK mice following SD relative to WT (Figure 4). Together, these findings support a role for *Arc* and its dependent IEGs in regulating rebound NREM sleep in conditions of forced wakefulness.

## Discussion

In this study, we use the Adk-Tg model to demonstrate distinct molecular pathways underpinning regulation of sleep need under baseline conditions and following sleep deprivation. We show that both sleep timing/distribution across the 24h day (correlating with Process C), and sleep amount (Process S) are altered when ADK is over-expressed. These changes correlate with distinct transcriptomic signatures in the SCN and cortex that are associated with altered synaptic signalling and changes in immediate early gene expression including of those that are known to regulate sleep (e.g., *Arc*). Furthermore, our data supports a role for ERK phosphorylation in inducing sleep and as a marker of sleep need under normal environmental conditions. We demonstrate that under SD however, ERK phosphorylation decreases and therefore is unlikely to drive rebound sleep, and the main signalling cascade associated with rebound NREM is likely the immediate early genes like *Arc*, *Fos*, *Nr4a1*, etc.

Under 12/12h LD cycles, Adk-Tg mice were awake longer, with more wake episodes through the light phase. These mice also showed an altered distribution of sleep at the light/dark transitions and a lower amplitude and more fragmented circadian rhythm (Figure 1). However, in constant dark, we see no difference during the subjective day, but the altered distribution at transitions persists (Figure 2). These data underscore that in addition to directly modulating Process S, adenosine signalling is an important determinant of Process C, the effect of light on Process C, and the interaction with Process S. The transcriptomic signatures of the SCN that may correlate with the changes in these processes could include more inhibitory signalling - increased GABA, decreased glutamate, and lowered expression of potassium channels - which are also reflected to some extent within the cortex. Whilst multiple neuropeptides mediate SCN cellular communication, the primary neurotransmitter of the SCN is GABA. As an inhibitory neurotransmitter, GABA is an important SCN output acting on distal GABAergic synaptic terminals and mediating dorsal-ventral communication within the SCN, and its extracellular concentration (maintained by rhythmic uptake by surrounding astrocytes) is crucial in SCN timekeeping (Brancaccio *et al*., 2017; Patton *et al*., 2023). Hence, an increase in the levels of the GABA_A_ subunit *Gabra2a* would suggest higher GABA sensitivity across the SCN and more inhibition. Furthermore, glutamate is the primary neurotransmitter mediating photic input to the SCN, and the alteration to the circadian rhythm under LD conditions may be related to this.

A prominent feature of the cortical transcriptome of the Adk-Tg is the increased mitochondrial activity at baseline (Figure 3), and the upregulation of oxidative phosphorylation and electron transport chain pathways following SD (Figure 4), presumably as a result of increased ADK function, with co-occurrence of attenuated DNA repair genes in both conditions. Indeed, wakefulness with active exploration promotes double stranded breaks in murine cerebral cortex, and lack of sleep impairs DNA repair (Bellesi *et al*., 2016). Hence, the overall reduction in NREM (Palchykova *et al*., 2010) and total baseline sleep (Figure 1) in Adk-Tg could be attenuating DNA repair. This is in line with other studies showing increased susceptibility to carcinogens in the Adk-Tg (El-Kharrag *et al*., 2019) and complex crosstalk at multiple levels between DNA damage and adenosine signalling, reviewed in (Stagg *et al*., 2023). IEGs like *Arc*, *Fos*, and *Npas4* are downregulated at baseline (Supplementary table 1) and following sleep deprivation these IEGs and markers of DNA damage like *Gadd45g* and *Parp1* are significantly upregulated (Figure 4C). Remarkably, *Gadd45g* has been shown to regulate IEG expression in murine cortex (Li *et al*., 2019), while the activity of the DNA damage detector *Parp1* has been shown to increase following SD and promote sleep (Zada *et al*., 2021). The increase in oxidative stress associated with ADK overexpression and the reduced baseline sleep duration could thus be exacerbating DNA-damage following SD, which is in part driving the increase in IEG expression and the increase in rebound NREM sleep, and this is independent of sleep pressure as SWA build up during SD in these mice is lower than their WT littermates (Palchykova *et al*., 2010). Hence this demonstrates the two-way relationship between sleep and DNA repair and stresses the importance of sleep, particularly NREM, in facilitating DNA repair following SD-induced damage. It also proposes a role for IEG induction in mediating rebound NREM.

Under basal conditions of normal light/dark cycles, ADK-Tg mice are awake longer (Figure 1) and exhibit lower pERK1/2 (Figure 5) compared to their littermate controls. Overexpression of ADK in mice leads to a significant reduction in overall NREM sleep (Palchykova *et al*., 2010). Similarly, previous reports have shown that pharmacological inhibition of ERK phosphorylation in mice increases wakefulness duration and lowers NREM sleep (Mikhail *et al*., 2017). However, the same study has demonstrated that ERK phosphorylation in the cortex accumulates during wakefulness, while global *Erk1* deletion or conditional knockdown of *Erk2* in cortical neurones both result in an increase in duration of wakefulness. Together these observations suggest that the build-up of pERK1/2 serves as a marker of sleep need under baseline conditions, and that the accumulation of activated pERK1/2 is needed to promote sleep. This is of particular importance, as it strengthens our understanding of ERK as a sleep-inducing kinase and a marker of sleep need under normal environmental conditions.

Following sleep deprivation however, adenosine and ERK signalling diverge. Similar to what has been previously reported (Guan *et al*., 2004; Vanderheyden *et al*., 2013; Mikhail *et al*., 2017), pERK1/2 levels decrease in both genotypes after 6 hours of sleep deprivation at the beginning of the light phase (Figure 5). Conversely, under similar conditions of sleep deprivation, ERK phosphorylation and adenosine signalling exhibit different rebound-sleep phenotypes (Palchykova *et al*., 2010; Mikhail *et al*., 2017). Pharmacological Inhibition of ERK phosphorylation is evidently accompanied by a significant decrease in NREM rebound sleep (Mikhail *et al*., 2017) while ADK-Tg mice have been shown to demonstrate a significant increase in NREM rebound sleep (Palchykova *et al*., 2010). This proposes that rebound NREM sleep is regulated by an ERK-independent pathway. On the other hand, the upregulation of IEGs, particularly *Arc*, could be the key driver for rebound NREM sleep following SD. Indeed, *Arc* is important for the induction of NREM and REM sleep, given *Arc* KO mice lack the rebound effects observed following sleep deprivation (Suzuki *et al*., 2020), and Adk-Tg mice show both increased *Arc* expression (Figure 4, Supplementary Figure 3) and increased rebound NREM sleep following SD (Palchykova *et al*., 2010). Furthermore, immediate early genes known to be under the direct effect of *Arc* (such as *Egr1*, and *Nr4a1*) are upregulated in ADK mice following SD relative to WT (Figure 4). Hence, it is possible that while the basal pERK1/2 and *Arc* expression are lower in ADK mice (contributing to lower sleep pressure and SWA build up), upregulation of *Arc* due to SD is significantly higher, which in turn increases rebound NREM sleep. And since *Arc* is associated with synaptic downregulation (Korb & Finkbeiner, 2011), together with the observed attenuation in synaptic signalling in Adk-Tg mice, these findings support the hypothesis that sleep, particularly NREM, reduces synaptic strengthening.

Interestingly, SRF was identified as a key transcription factor involved in the differential gene expression observed in Adk-Tg mice under baseline conditions (Figure 3) and has been shown to regulate expression of *Arc* in neurones (Pintchovski *et al*., 2009). In addition, SRF activity correlates with sleep pressure (Hor *et al*., 2019) and its reduction increases DNA stress (Zhang *et al*., 2023). *Arc* being downregulated in Adk-Tg mice suggests a reduction in SRF transcription factor activity. Therefore, under baseline conditions, Adk-Tg mice display reduced SRF-dependent transcription, reduced pERK1/2 levels, and less SWA build up (Palchykova *et al*., 2010), emphasizing that all of the above correlate with reduced sleep pressure and baseline sleep. On the other hand, under forced wakefulness where the homeostatic arm of Process S is involved, Adk-Tg display an increase in rebound NREM which is independent of sleep pressure, since SWA build up during SD in these mice is lower than in their WT controls (Palchykova *et al*., 2010), but mainly correlates with the upregulation of IEGs (Figure 4). Together, this proposes that baseline and homeostatic components of process S could be governed by different mechanisms.

In conclusion, our results suggest adenosine signalling plays an important role in integrating Process S and Process C, but that distinct molecular mechanisms control baseline sleep amount versus rebound sleep.

## EXPERIMENTAL MODELS

### Animals

C57BL/6J mice and Adk-Tg transgenic mice (Fedele *et al*., 2005) of both sexes were used, typically aged 80 days to 6 months. All procedures were performed in accordance with the UK Home Office Animals (Scientific Procedures) Act 1986 and the University of Oxford’s Policy on the Use of Animals in Scientific Research (PPL 70/6382, PPL 8092CED3), as approved by the local Animal Care and Ethical Review committee (ACER). Animals were sacrificed via Schedule 1 methods in accordance with the UK Home Office Animals (Scientific Procedures) Act 1986.

## METHOD DETAILS

### Behavioural assay

Adk-Tg (n=10) and wildtype mice (n=10) were singly housed, passive infrared activity recorded, and light/dark cycles modified as indicated on the figures (light levels approx. 200 lux from white LED). Data were analysed on Clocklab (Actimetrics, Wilmette, IL). Immobility defined sleep was analysed as reported in Fisher et al (Fisher *et al*., 2012).

### Sleep deprivation

In independent experiments, C57BL/6 (n=8) and Adk-Tg (n=6) mice between 8-12 weeks of age were housed under a 12-h light/dark cycle with *ad libitum* access to food and water. Mice of the same sex were group-housed in the same cage. The sleep deprivation experiment was conducted on the first day after transfer to DD. The protocol consisted of gentle handling and novel object introduction between CT0-6 under dim red light. Sham-treated animals were allowed to sleep in constant dark. At the end of the protocol, both sleep-deprived and sham-treated mice were sacrificed by cervical dislocation under dim red light.

### Tissue collection

#### Tissue collection for mRNA analysis

After animal sacrifice, eyes were removed before lights were turned on. Brains were rapidly extracted, placed into a brain matrix (Kent Scientific, Torrington CT, USA) and kept ice-cold. To collect the cortex, a 1mm thick coronal section was dissected using skin graft blades (Swann-Morton, Sheffield, UK) positioned at Bregma -0.10 mm and -1.10. A biopsy punch (1mm diameter – Integra Life Sciences, York, USA) was used to collect SCN and cortex tissue punches which were frozen on dry ice and stored at −80 °C before RNA extraction took place.

### *In vitro* experiments

#### RNA extraction

Total RNA was extracted according to a modified version of the RNeasy Micro Kit (Qiagen, Germany). Tissues were first homogenized in 100 µL of TRIzol Reagent (Thermo Fisher Scientific, USA). Then, 400 µL of TRIzol and 100 µL of chloroform were added and samples thoroughly mixed. The samples were then centrifuged for 15 minutes at 4°C. The resulting supernatant was collected, and RNA was extracted based on RNeasy Micro Kit’s RNA extraction protocol. RNA quality and quantity was checked using Nanodrop1000 and Qubit RNA High Sensitivity, Broad Range Assay Kits (Thermo Fisher Scientific, USA).

#### RNA-seq library preparation

100 ng of RNA was used to prepare libraries using the Illumina Stranded Total RNA Prep, Ligation with Ribo-Zero Plus kit following the manufacturer’s instructions. Following library preparation, library concentration was determined using the Library Quantification Kit for Illumina Platforms (Roche, UK) following the manufacturer’s instructions. RNA-seq processing was performed by Novogene (Cambridge, UK).

#### RNA-seq data analysis Data pre-processing

All computational analyses were performed using a combination of local machines and the Biomedical Research Council’s high-performance computing cluster. >10 million 150-bp paired-end reads were obtained for each replicate. The quality of raw sequencing data was assessed using FastQC v0.11.9 (Andrews, 2010) both before and after adapter trimming. All samples showed passing scores on all FastQC criteria. Nextera adapter sequences, low-quality base calls (Phred score < 15) and short reads (read length < 20 base pairs) were trimmed using Trim Galore v0.6.2 (Martin, 2011). FASTQ paired-end reads were aligned using HISAT2 v2.2.1 (Kim *et al*., 2019) to mouse GRCm38 genome build from Ensembl (Zerbino *et al*., 2018). BAM files were sorted by read name and chromosome position using Samtools v1.8 (Li *et al*., 2009). Transcripts were quantified via the FeatureCounts function of Rsubread v1.6.4 (Liao *et al*., 2014).

#### Differential analysis

For comparison between sleep-deprived and sham-treated samples, downstream differential analysis was performed on the count matrix using R version 4.2 and R package DESeq2 v3.15 (Love *et al*., 2014). Data was normalized using DESeq2’s built-in median-of-ratios method to account for library depth and RNA composition across samples. Genes with low counts (sum of counts is less than 30 across all samples) were filtered, as they mostly reflect noise in the dataset. For comparison of the transcriptomic response to sleep deprivation between the C57BL/6 and Adk-Tg genotypes, to account for baseline differences in transcript quantity, DESeq2’s ratio-of-ratio method was used. P values were adjusted using BH method for multiple testing. Significant differentially expressed genes (DEGs) were selected with an adjusted p-value < 0.1 for comparison between sleep-deprived and control samples and adjusted p-value < 0.1 and log_2_ fold change ≥ 0.5 or ≤ -0.5 for comparison between genotypes.

#### Pathway enrichment analysis

Significant DEGs were used to identify over-represented Gene Ontology (GO) (Carbon *et al*., 2019) terms (GO Biological Process and Molecular Function databases) and enriched pathways annotated in the Kyoto encyclopedia of genes and genomes (KEGG) database (Kanehisa *et al*., 2016) using the R package gprofiler2. Protein-protein interaction of the gene modules was constructed using the STRING protein database (https://string-db.org/) and visualized on CytoScape 3.8.

#### Western Blotting

Twenty μg of brain tissue total protein in RIPA buffer were run on 10% SDS-PAGE gels (Novex, Life Technologies), transferred using standard protocols (Bio-Rad) onto Transblot Turbo LF PVDF membranes (Bio-Rad Laboratories), blocked with Odyssey blocking buffer (Li-Cor Biosciences, Lincoln, NE, USA) incubated with primary antibodies to ERK1/2 (Rabbit anti- p44/42 MAPK ERK1/2 137F5, #4695 Cell Signalling Technology used at 1:1000), pERK (Rabbit anti- p44/42 MAPK ERK1/2 T202/204, #4370 Cell Signalling Technology, used at 1:1000), and B-actin (Mouse anti-B-actin, #66009-1-lg Proteintech, used at 1:20,000), subsequently with IRDye 680 donkey anti-rabbit IgG or IRDye 800 goat anti-mouse secondary antibody and scanned with the Odessey Li-Cor system.

#### qRT-PCR validation

cDNA was synthesized from the purified RNA according to the qScript cDNA Synthesis Kit (Quanta Biosciences). mRNA was quantified using the QuantiFast SYBR® Green PCR Kit (QIAGEN) in a StepOnePlusTM thermal cycler (Applied Biosystems).

Cycle conditions were 95 °C for 5 min, and 40 cycles of 95 °C for 10 s, 60 °C for 30 s, 72 °C for 12 s. The cycle thresholds for each gene were normalized to the geometric mean of *Rps9* and *B2m* housekeeping genes. The relative gene expression between SD and control samples was calculated using the 2 −ΔCt method. To ensure that punches of the defined areas were accurately dissected and that the animals were indeed sleep deprived, qPCR of the cortex-specific mRNA *Arc* and *Nr4a1* were conducted (Thompson *et al*., 2010b), revealing selective enrichment. Primer sequences are listed below.

### Primer sequences

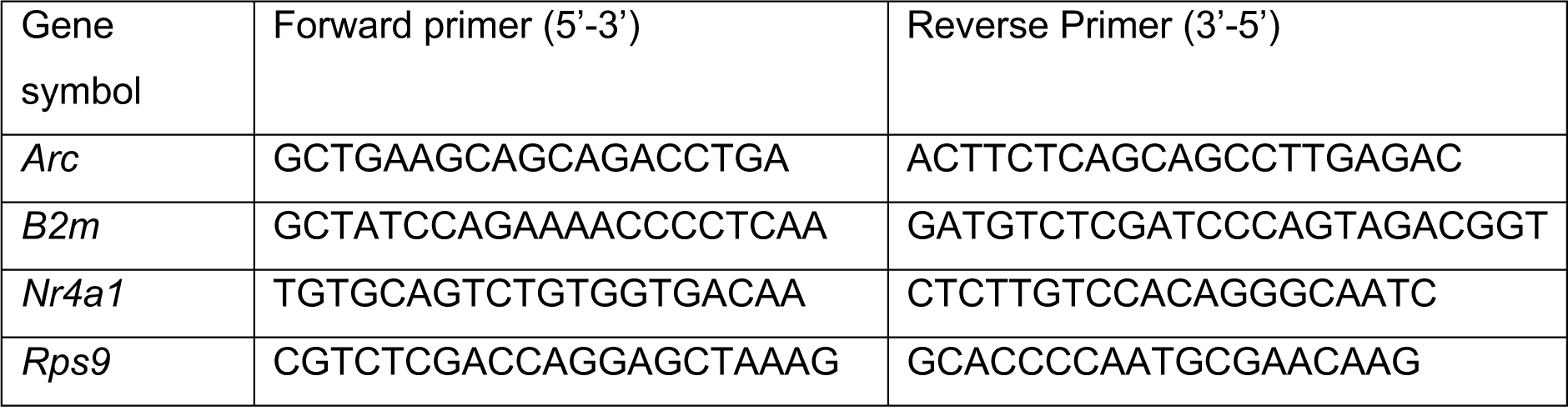

### Statistical analysis

N represents the number of independent animals or replicates per group, as detailed in each figure legend. To correct for errors of multiple testing, the false discovery rate (FDR) was calculated for each statistical test. The Benjamini and Hochberg (BH) method was used to compute the false discovery rate (FDR) for each statistical test. The thresholds for DEG detection were specified in the Methods section. Statistical testing was performed using built-in packages in R (version 4.2) (R Core Team, 2022) and Python (version 3.10) (Van Rossum & Drake Jr, 1995).

## Acknowledgements

This work was supported by the following sources of funding: BB/N01992X/1 David Phillips fellowship from the BBSRC to AJ, Elysium-Oxford Post-Doctoral Fellowship to LT. We would like to thank Prof. Steven Brown for insightful discussions and for generously providing behavioural data pertaining to the *Adk-Tg* mice to Prof. Dallmann prior to his untimely passing.

## Data accessibility Statement

All data will be deposited at NCBI SRA upon acceptance

---

**Supplementary Figure 1:**
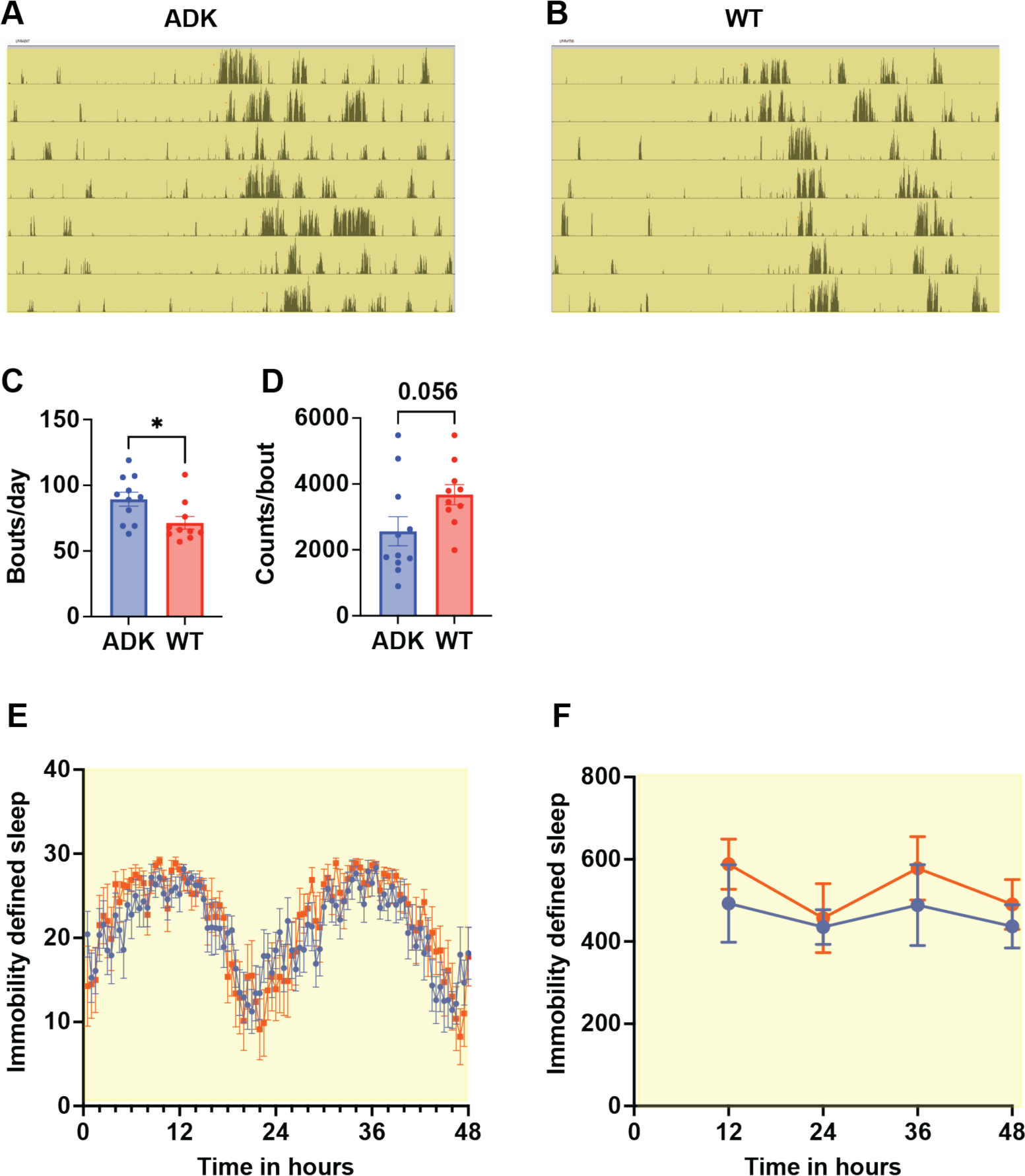
Adk-Tg mice show altered distribution of sleep and increased fragmentation under constant light. Representative actograms from (A) Adk-Tg (ADK) and (B) wildtype (WT) mice under constant light, yellow indicates lights on and black bars are passive infrared indicated activity. Periodogram analysis showed significant differences between the genotypes in (C) number of bouts per day (D) the activity counts/bout. No clear differences in immobility defined sleep distribution (in minutes) across 2 days in LL were observed (E) but overall differences (F) in total sleep levels were seen (p=0.002, (F(1, 72) = 15.18) for genotype effect with two-way ANOVA analysis).

**Supplementary Figure 2:**
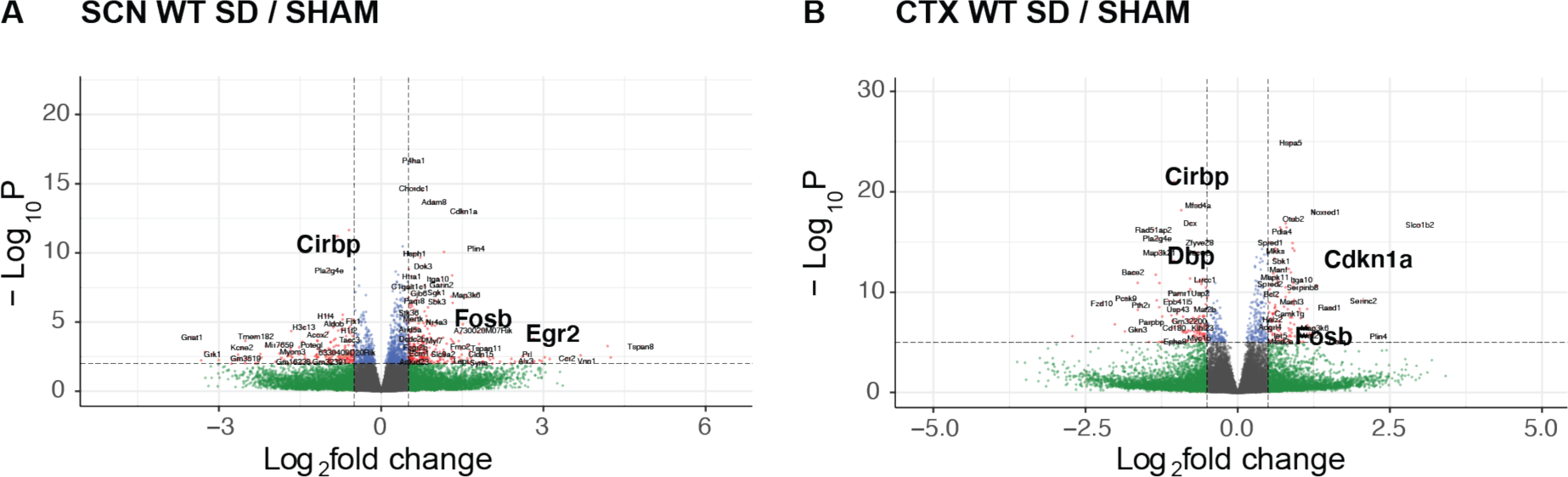
Wildtype mice show transcriptomic changes in SCN and cortex in response to sleep deprivation. Volcano plots of RNA-Seq data from (A) SCN and (B) Cortex (CTX) of wildtype (WT) mice sleep deprived (SD) for 6h (CT0-6) versus sham deprived (SHAM) mice. Dotted line horizontal indicates adjusted -Log_10_ P = 2 (SCN) and 5 (CTX), dotted vertical lines indicate Log_2_ fold change of ±0.5. Green = transcripts with Log_2_ fold change > ±0.5, red = transcripts with Log_2_ fold change > ±0.5 and adjusted p<0.01 (SCN) and p < 0.0001 (CTX).

**Supplementary Figure 3:**
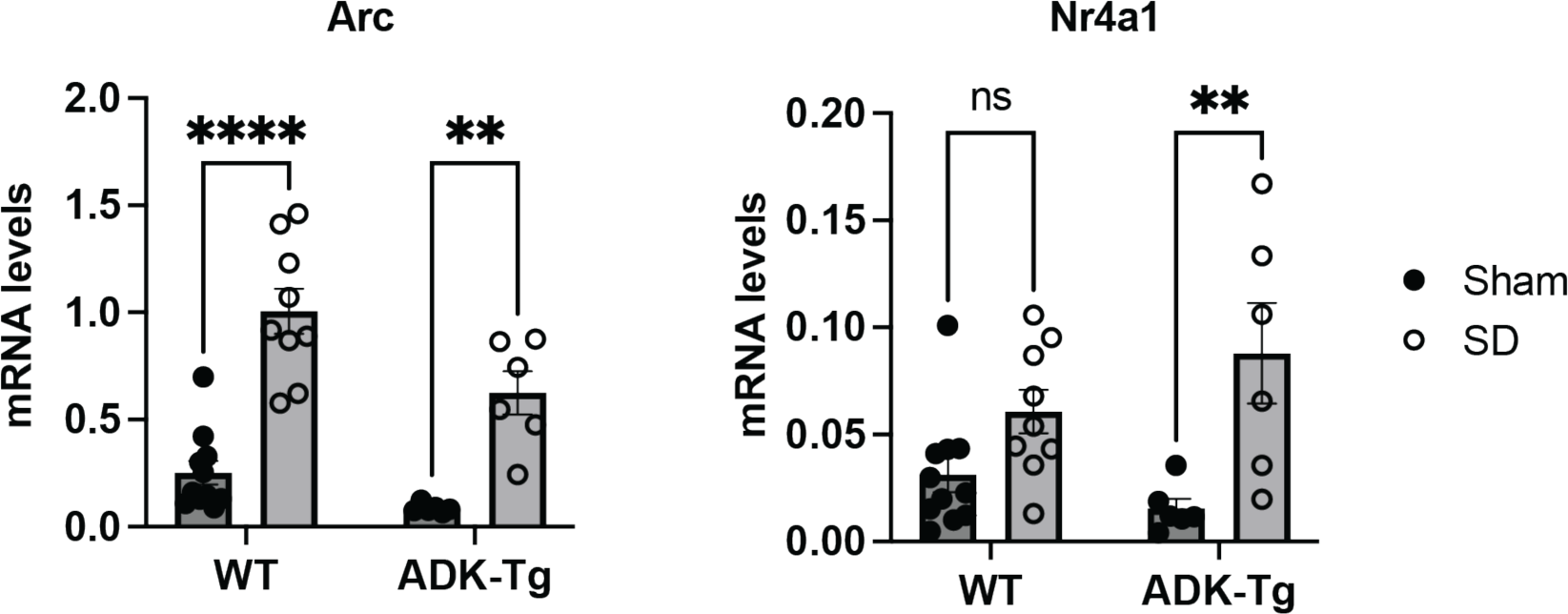
qPCR validation of *Arc* and *Nr4a1* changes in the cortex in response to sleep deprivation. Adk-Tg and wildtype (WT) mice sleep deprived (SD) for 6h (CT0-6) versus sham deprived (Sham) mice, mRNA levels of transcripts shown relative to housekeeping genes (see methods). ** = p<0.01, **** = p < 0.0001, student’s t-test, n = 6-8.

